# Multi-region proteome analysis quantifies spatial heterogeneity of prostate tissue biomarkers

**DOI:** 10.1101/250167

**Authors:** Tiannan Guo, Li Li, Qing Zhong, Niels J Rupp, Konstantina Charmpi, Christie E Wong, Ulrich Wagner, Jan H Rueschoff, Wolfram Jochum, Christian Fankhauser, Karim Saba, Cedric Poyet, Peter J Wild, Ruedi Aebersold, Andreas Beyer

**Author notes:** Equal contribution (Tiannan Guo, Li Li, Qing Zhong). Corresponding authors Email contacts for corresponding authors: Peter J. Wild Ruedi Aebersold Andreas Beyer.

## Abstract

Many tumors are characterized by large genomic heterogeneity and it remains unclear to what extent this impacts on protein biomarker discovery. Here, we quantified proteome intra-tissue heterogeneity (ITH) based on a multi-region analysis of 30 biopsy-scale prostate tissues using pressure cycling technology and SWATH mass spectrometry. We quantified 8,248 proteins and analyzed the ITH of 3,700 proteins. The level of ITH varied significantly depending on proteins and tissue types. Benign tissues exhibited generally more complex ITH patterns than malignant tissues. Spatial variability of ten prostate biomarkers was further validated by immunohistochemistry in an independent cohort (n=83) using tissue microarrays. PSA was preferentially variable in benign prostatic hyperplasia, while GDF15 substantially varied in prostate adenocarcinomas. Further, we found that DNA repair pathways exhibited a high degree of variability in tumorous tissues, which may contribute to the genetic heterogeneity of tumors. This study conceptually adds a new perspective to protein biomarker discovery by quantifying spatial proteome variation and it demonstrates the feasibility by exploiting recent technological progress.

## Introduction

During the last decade numerous new cancer treatment options have been developed. Their optimal application, however, requires better molecular characterization of the tumors with the aim of developing biomarkers matching the specific tumor to the best available therapy. Some cancer types, such as prostate cancer, still suffer from an ‘over treatment problem’, *i.e.* radical therapy such as removal of the organ in unnecessary cases due to uncertain diagnosis. These problems persist despite the recent progress in genomic, transcriptomic, and proteomic profiling of tumors. In contrast to the standardization of histopathological diagnostic categories, tumor grading, and standards of reporting, molecular testing is still underexploited in routine diagnostics of localized prostate cancer cases. A recent review about biomarkers in prostate cancer (Kristiansen, 2018) has highlighted the need to consider intra-tissue heterogeneity (ITH) in each individual case for successful molecular testing. ITH is of high clinical relevance. For instance, a tumor may contain a small sub-population of cells with primary resistance, leading to incomplete response to treatment or early recurrence (Murtaza, Dawson et al., 2015). High degree of Gleason score, DNA ploidy, and PTEN expression have been observed in prostate tumors (Cyll, Ersvaer et al., 2017). Thus, it remains a challenge to optimize clinical decisions based on single biopsies (Boutros, Fraser et al., 2015).

Indeed, ITH is an important contributor to spatially variable molecular levels, which poses a substantial problem for biopsy-based tumor diagnostics, because for highly variable proteins, the measured quantity is position-dependent. Genomic ITH has been predicted based on clonal evolution and the cancer stem cell hypothesis (Dalerba, Cho et al., 2007). This prediction was experimentally validated by the application of high-throughput sequencing to small tissue samples and even single cells. Such studies have uncovered a high degree of genetic ITH in colon (Jones, Chen et al., 2008), pancreas (Yachida, Jones et al., 2010), breast (Russnes, Navin et al., 2011), prostate (Haffner, Mosbruger et al., 2013), renal carcinomas (Gerlinger, Rowan et al., 2012), and leukemia (Cancer Genome Atlas Research, 2013, Ding, Ley et al., 2012), with regard to both mutational and gene expression profiles of tumor cells. For example, Boutros *et al.* observed extensive ITH in prostate cancers at the level of gene copy number alterations and point mutations, which led to spatially divergent mutational patterns for thousands of genes, including several tumor-relevant genes (Boutros et al., 2015). It can be expected that genomic ITH will be translated, at least to some extent, to ITH at the protein level. For example, androgen receptor and prostate specific antigen (PSA/KLK3) expression can significantly vary between different regions within the same prostate carcinoma (Magi-Galluzzi, Xu et al., 1997, Shah, Bentley et al., 2015). Thus, there is a need to systematically describe and quantify protein level heterogeneity in tumor tissues.

Despite this well recognized need, technical challenges have so far prevented the quantification of protein level heterogeneity in tumor specimens at the proteomic scale (Alizadeh, Aranda et al., 2015). High-throughput antibody-based immunohistochemistry staining has been applied to tissue sections (Uhlen, Fagerberg et al., 2015). However, such data are semi-quantitative and limited in scope by the availability of suitable antibodies. Label-free shotgun proteomics has been used to explore in-depth the proteome of multiple regions of tumor tissues (Wisniewski, Ostasiewicz et al., 2012). However, due to the inherent technical limitations, the method is not suitable to systematically explore ITH at high sample-throughput and high spatial resolution, which is essential to achieve adequate spatial resolution (Domon & Aebersold, 2010). Single-cell proteomics using mass cytometry is another promising technology allowing quantification of protein levels in thousands of individual cells. However, the technique at present only measures 10s of proteins per sample (Giesen, Wang et al., 2014).

We have recently developed a mass spectrometry-based proteomics method, *i.e.* pressure cycling technology and sequential windowed acquisition of all theoretical fragment ion mass spectra (PCT-SWATH)(Guo, Kouvonen et al., 2015b), which supports highly reproducible and accurate quantification of a few thousand proteins from biopsy-scale tissue samples at high throughput. This is accomplished by the integration into a single platform of optimized sample preparation, mass spectrometric and computational elements. To generate mass spectrometry-ready peptide samples from tissue samples we adopted PCT to lyse the tissues, extract proteins and digest them into peptides in a single tube under precisely controlled conditions (Powell, Lazarev et al., 2012). To analyze the resulting peptide samples, we used SWATH-MS, a massively parallel targeting mass spectrometry method (Gillet, Navarro et al., 2012). In SWATH-MS all MS-measurable peptides in a sample are fragmented and periodically recorded over a single dimension of relatively short chromatography (Gillet et al., 2012). The net result of this technique is a single digital file that contains fragment ions of all mass spectrometry-detectable peptides, from which peptides and proteins are identified and quantified post acquisition, via a targeted data analysis strategy (Gillet et al., 2012, Röst, Rosenberger et al., 2014).

In this study, we approached proteomic ITH for prostate cancer tissues by PCT-SWATH-based multi-region proteomic analysis of 60 biopsy-level tissue samples from three prostate cancer patients. We then computed the technical and spatial biological variation for each measured protein in different types of tissues and different patients, and established a proteome-scale landscape of protein ITH in benign and malignant prostate tissues. Our data revealed distinct ITH patterns of prostate cancer biomarkers that were further independently validated using immunohistochemistry (IHC) in an independent set of 83 patients.

## RESULTS

### Study design for quantifying proteomic variability

We designed a study to quantify spatial proteomic variability in multiple regions of malignant and matching benign prostate tissues using the PCT-SWATH-MS platform (Guo, Kouvonen et al., 2015a). We assumed that the total proteomic variability observed in the sample cohort was composed of technical and biological variation, the latter including inter-patient, inter-tissue and intra-tissue variation. To open the possibility to partition the overall observed variability into its possible sources, we obtained tissue samples from multiple regions of prostatectomy specimens as illustrated in **Figure 1**. Each sample was a tissue punch biopsy consisting of a cylinder of 1 mm diameter and about 3 mm length that was derived from fresh frozen tissue blocks using a core needle. Samples were obtained from prostatectomy specimens in three individuals diagnosed with adenocarcinoma (ADCA) of the prostate. Gleason grading was performed according to the International Society of Urological Pathology and the World Health Organization consensus (Epstein, Egevad et al., 2016, Humphrey, Moch et al., 2016) (**Supplementary Fig. 1**). In total, 12 benign prostatic hyperplasia (BPH) and 18 ADAC tissue samples were obtained. One of the three individuals had a mixed acinar and ductal ADAC, and both subtypes were included in the study to measure the variation resulting from morphologically distinct subtypes. The other two patient samples displayed acinar ADCA by histologic means. Each tissue type (malignant *versus* benign) of each patient was sampled three to six times resulting in a total of 30 biological samples. Each sample was processed by PCT-SWATH in duplicate to evaluate the technical variation of the proteomic analysis (**Fig. 1, Supplementary Table 1**). The samples were grouped into 10 batches of six samples, according to patient identity, tissue type and technical replicate (**Fig. 1A, Supplementary Table 2**). This experimental design allowed us to subsequently estimate intra-tissue variability from within-batch comparisons (see **Methods**), which is important to avoid overestimating variances due to batch effects.

**Figure 1.**
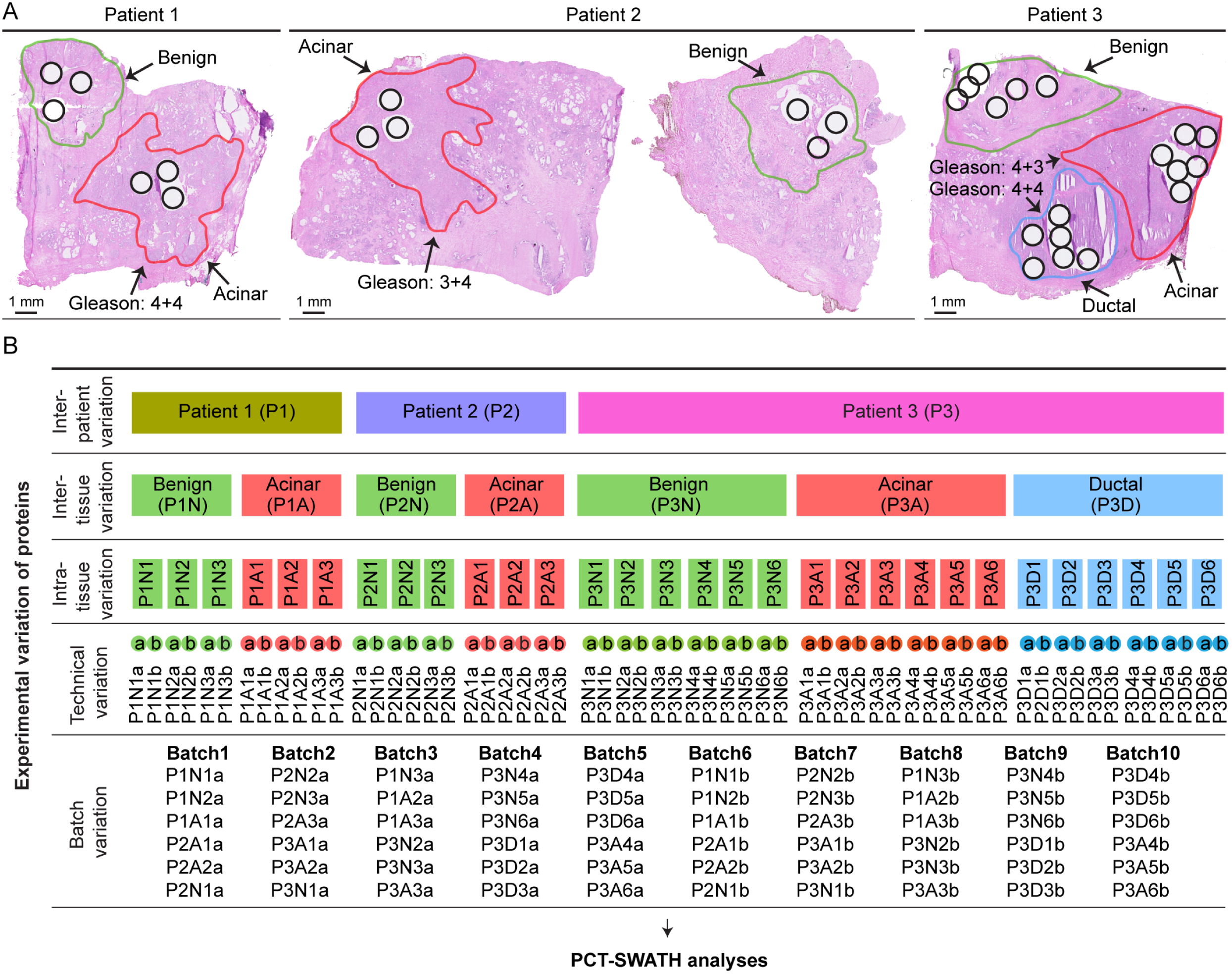
**Study design.** (**A**) H&E staining of the fresh frozen prostate tissue from three individuals who have contributed BPH (non-tumorous) and matching acinar or ductal adenocarcinoma. Green, orange, and blue lines depict regions diagnosed by a pathologist as BPH, acinar and ductal tumors, respectively. Black circles indicate where the punches were made. (**B**) Overall measured variation of protein expression was partitioned into biological and technical variation including inter-patient variation, inter-tissue variation, intra-tissue variation and technical variation from MS analysis and batch variation. Three or six punches were sampled from each tissue type, followed by PCT-SWATH analyses in technical duplicate. The samples were shuffled and analyzed in 10 batches of six samples.

### Quantitative proteomic analysis of 30 prostate cancer tissue regions

The 10 batches of samples were processed using PCT-SWATH in duplicate over a period of 15 working days. The acquired SWATH-MS data were subjected to *in silico* targeted analysis using the OpenSWATH software(Röst et al., 2014). In total, 39,493 proteotypic peptides and 8,248 protein groups were quantified consistently across all 60 measurements (**Supplementary Table 3 and 4**). The measured protein intensities were highly reproducible (average Pearson correlation values between replicates: 0.944). To obtain high-confidence estimates of ITH, we subsequently narrowed our statistical variation analyses to a subset of 3,700 proteins quantified by at least two concordant proteotypic peptides. Our peptide selection procedure ensured that the selected peptides showed consistent behavior across samples. Thereby, we minimized the possibility that peptide intensity variation was not due to protein abundance changes, but due to post-translational modifications or other artifacts (see **Methods**) (Picotti, Clement-Ziza et al., 2013). We then corrected batch effects in the dataset by subtracting the average signal of each protein per batch. After batch correction, most technical replicates grouped together by unsupervised clustering based on the abundance of all proteins (**Supplementary Fig. 2**).

### Quantification of spatial proteomic heterogeneity

Our estimates of proteomic ITH are based on the notion that the signal variation between two samples is due to a combination of biological and technical factors. Since the biological variation is not directly quantifiable, we estimated biological variance by subtracting the technical variance from the total observed punch-to-punch variance.

The technical variance was estimated by calculating the dispersion between two technical replicates for each sample (independent protein digests from the same punch measured separately), *i.e.* generating 30 initial technical variance estimates per protein before averaging them (see **Methods** for details). This strategy produced seven technical variance estimates for all pairs of patient / tissue type (three normal tissue regions, three acinar tissue regions, and one ductal tissue region, **Fig. 1**). Pairwise correlations of these seven independent estimates showed that technical variances were consistently positively correlated, with a median correlation of 0.572 (**Fig. 2A**). Likewise, we analyzed the same type of correlation for the total punch variances. Like the technical variance, independent estimates of the total variance were also highly correlated, albeit with a slightly lower median correlation of 0.302, suggesting that the technical variance was more robust and less dependent on the specific sample than the total variance and the biological variance (**Fig. 2B**). Thus, as expected, the technical variance of a protein was mostly determined by its physico-chemical properties, whereas total variance varied in different tissue samples probably due to biological factors. Further, technical variance of log-transformed intensities was independent of the mean log-intensity **(Supplementary Fig. 3)**, suggesting that the same estimate of technical variance could be used at high and low protein concentrations. Subsequently, we averaged the seven estimates of technical variance per protein to obtain a single, robust estimate of each protein’s technical variance.

**Figure 2.**
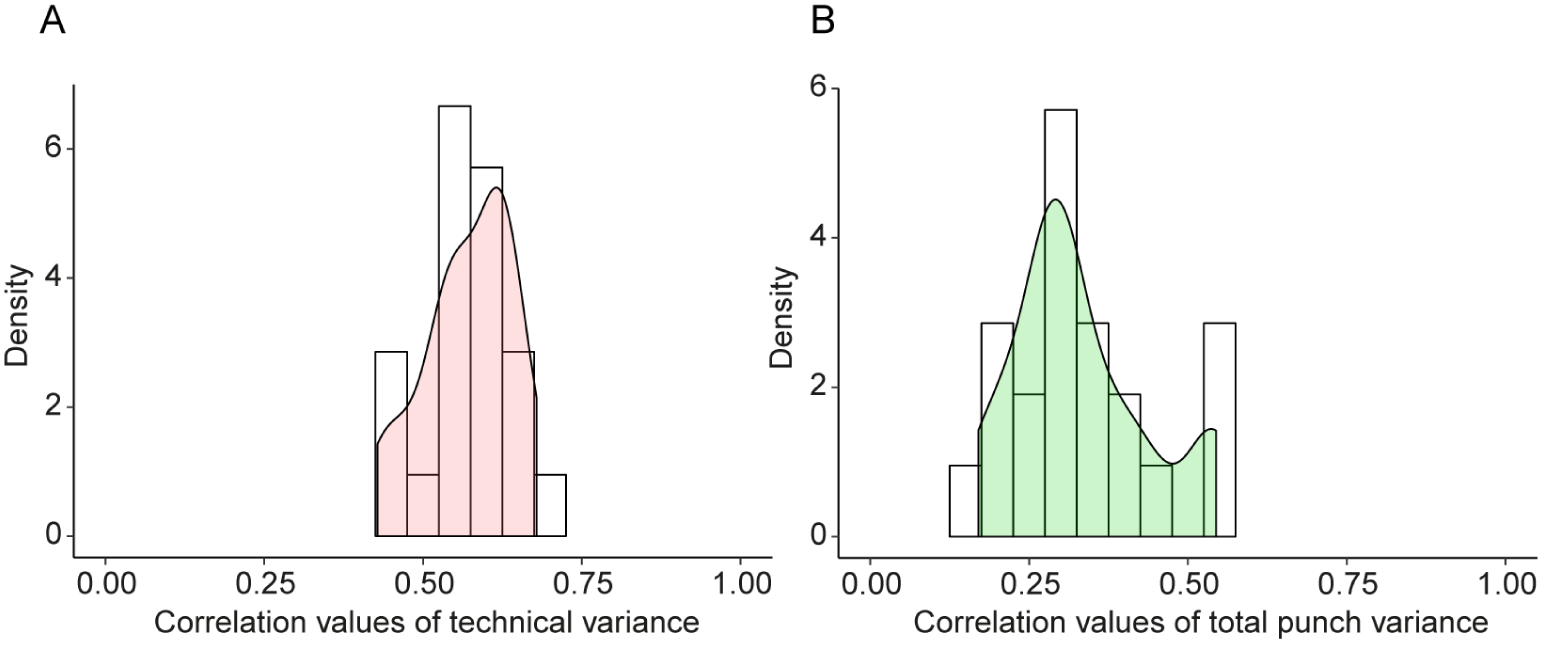
**Consistency of technical and total variance. (A)** Correlation of technical variances estimated independently for different samples. Technical variance is estimated from technical replicates. **(B)** Correlation of total variances (between punches) estimated independently from punches from different tissue samples (different patients, different tissue types).

Having established that our estimates of total variances and technical variances are robust, we next computed biological variances by subtracting each protein’s technical variance from its total variance between punches of the same patient and tissue type (see **Methods**). This yielded an estimate of intra-tissue biological variances of protein abundance which can be interpreted as the degree of proteomic ITH. The technical and total variances were independently estimated, which makes it numerically possible that the technical variance can be larger than the total variance of a specific set of punches. Indeed, for 183 proteins (4.9%) the estimated technical variance was larger than the total variance (**Supplementary Fig. 4**). These were mostly the proteins with very low total variance. We could not rigorously quantify the biological variances of these proteins, nevertheless, we assumed that most of them would have comparably low biological variances. Proteins with technical variances higher than total variances were excluded from most subsequent analyses.

Next, we compared the biological variances within a tissue with the biological variance between tissue types (benign versus malignant; termed *inter-tissue*) and between patients (**Fig. 3**). Inter-tissue and inter-patient variances were obtained by first averaging protein intensities from punches of the same tissue or patient, respectively (see **Fig. 1A** and **Methods**). Our data showed that the biological variance between punches within the same tissue (*i.e*. intra-tissue variance) is of similar magnitude as the variation of average intensities between tissues and patients, indicating a high degree of protein ITH (**Fig. 3A**). Further, the protein variances between patients, tissue, and within tissue were significantly correlated (**Fig. 3B-D**). Thus, a protein with large intra-tissue variation is also likely to vary across tissues and between the three patients.

**Figure 3.**
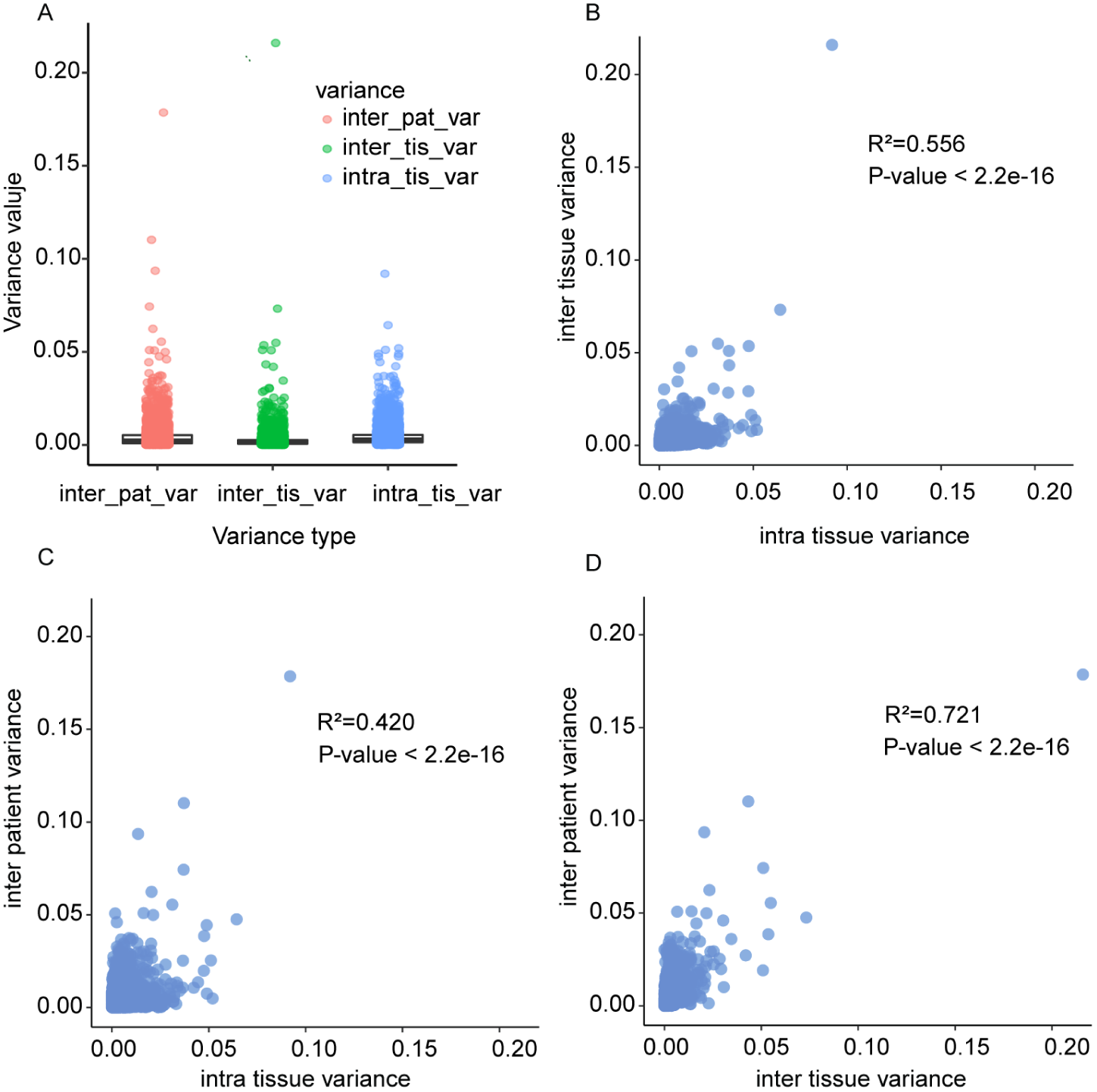
**Correlation of biological variance between patients and tissue types.** Each dot represents one protein. **(A)** Distributions of biological variance estimates. Inter-patient variances and inter-tissue variances are based on averaging the measurements of at least three punches. Intra-tissue variance was first determined independently per patient and tissue type, and then averaged. **(B)** Biological variance between tissue of the same patient versus variance between punches of the same patient and tissue. **(C)** Biological variance between different patients but same tissue type versus variance between punches of the same patient and tissue. **(D)** Biological variance between the same tissue types in different patients versus variance between different tissue types of the same patient.

### Classification of proteins based on their intra-tissue variability

To characterize ITH in different tissue types, we compared the biological variance of each protein in benign and malignant prostate tissues, and quantified the variability of 3,517 proteins in BPH and ADCA tissue samples (**Supplementary Table 5**). Interestingly, we observed a strong dependence of the variability of some proteins on the tissue type. We then classified the thus quantified proteins into five groups based on their biological variance patterns in the different sample types (**Fig. 4A**). Group no. 1 consisted of 100 proteins that were always robust and generally showed little intra-tissue variation in benign and malignant prostate tissues. Group no. 2 consisted of 339 proteins that varied substantially more in benign tissues compared to malignant tissues. Group no. 3 consisted of 93 proteins that varied more strongly in malignant tissues compared to benign tissues. Group no. 4 contained 365 proteins that had high intra-tissue variance in both malignant and benign tissues, while group no. 5 contained the remaining 2,620 proteins with intermediate variability. Remarkably, the top three most variable proteins in BPH are three proteins known or used in the diagnosis of prostate tumors, including prostate-specific antigen (PSA/KLK3), prostatic acid phosphatase (PAP/ACPP) and Desmin (DES). PSA is an androgen-regulated kallikrein family serine protease, that is produced by the secretary epithelial cells in acini and ducts of prostate glands (Balk, Ko et al., 2003). The secreted PSA, originated from prostate tissues, is the most commonly used, blood-based biomarker for prostate cancer (Hayes & Barry, 2014). However, PSA screening has remained controversial because of uncertainty surrounding its benefits and risks and the optimal screening strategy (Barry, 2009). Our data showed that PSA *in situ* was most variable in BPH but more stable in ADCA tissues. Since PSA is regulated by androgen, this indicates androgen-driven malignant growth of prostate tumor cells. PAP is a non-specific tyrosine phosphatase and a well-studied tumor suppressor for PCa. PAP has already been used in immunotherapy regimens against PCa (Di Lorenzo, Buonerba et al., 2011) and is the second most variable protein in BPH after PSA. The variability of PAP expression was relatively high in ADCA samples, but lower than its variability in BPH samples. Desmin (DES) constructs class-III intermediate filament in smooth muscle cells. As a marker for prostate stromal composition, DES expression has already been associated with PCa survival (Ayala, Tuxhorn et al., 2003). Tuxhorn et al. have shown that prostate cancer-reactive stroma is composed of a myofibroblast/fibroblast mix with a significant decrease or complete loss of fully differentiated smooth muscle, whereas normal prostate stroma is predominantly smooth muscle (Tuxhorn, Ayala et al., 2002). Given the known heterogeneous composition of myoglandular hyperplasia (*i.e.* BPH) out of glandular and stromal (smooth muscle) elements, the higher variability of DES expression in BPH compared to PCa is not surprising.

**Figure 4.**
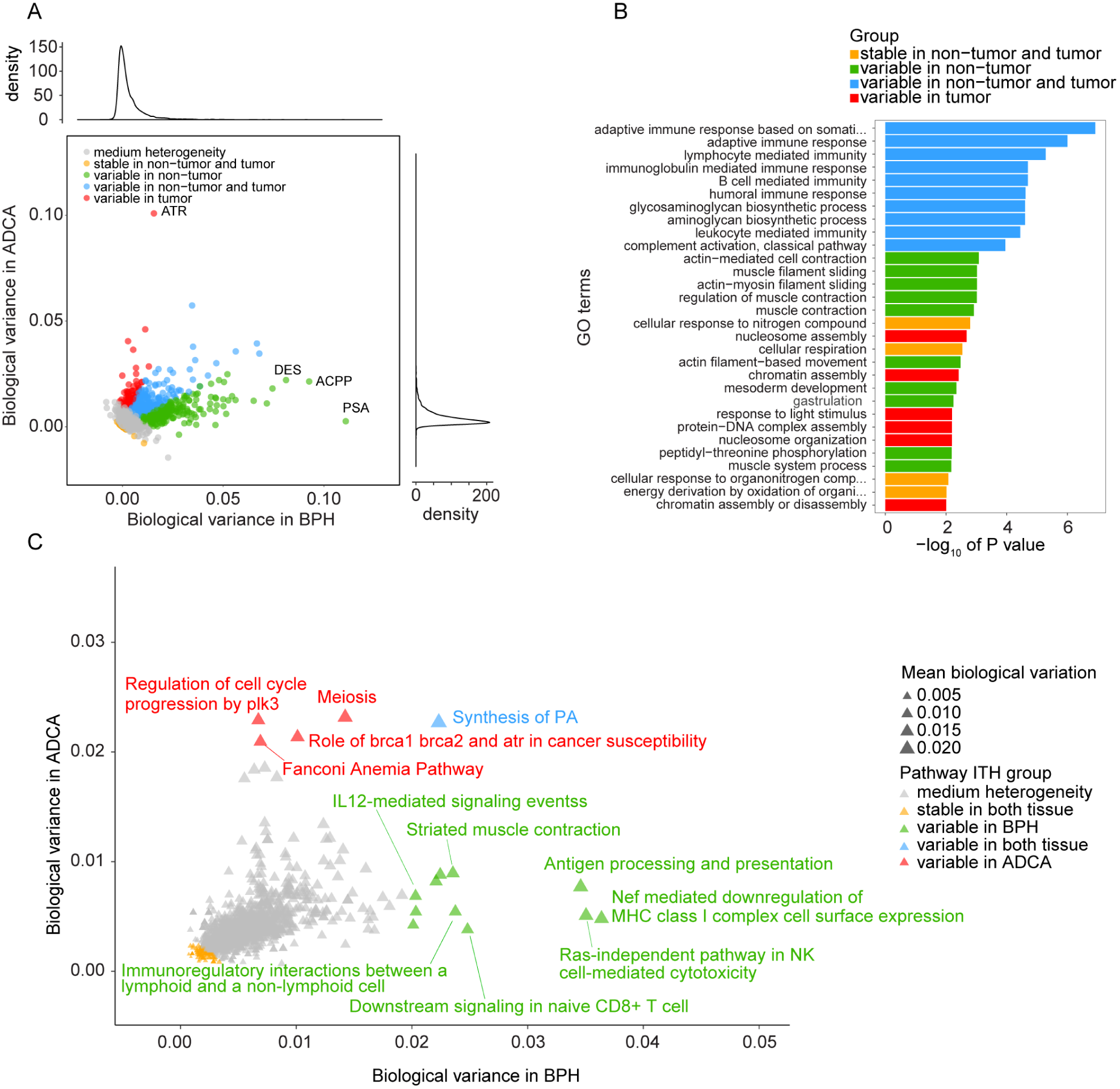
**Intra-tissue heterogeneity in tumorous and non-tumorous tissue.** (**A**) Biological variance among punches from the same tissue region was considered as the degree of intra-tissue heterogeneity for the respective tissue type. Degree of intra-tissue heterogeneity for each protein in benign versus malignant tissue are shown and colored according to classification. (**B**) GO enrichment analysis of four protein categories from (**A**). Length of horizontal bars indicates the significance of the enrichment. (**C**) Intra-tissue heterogeneity of biochemical pathways. Each triangle is the average biological variance (intra-tissue heterogeneity) of all quantified proteins from the respective pathway. Degree of intra-tissue heterogeneity for each pathway in benign versus malignant tissue are shown. Pathways were grouped according to their variability in benign and malignant tissue.

To further investigate the protein variability classes, we then performed a gene ontology (GO) enrichment analysis (**Fig. 4B**). As expected, stable proteins of group no. 1 were enriched for basic cellular functions that were required irrespective of the tissue state, such as energy metabolism (**Fig. 4B**). Proteins highly variable in both malignant and benign tissues (group no. 4) were enriched for immunity-associated processes. Muscle-related proteins exhibited a high degree of heterogeneity in benign tissues, reflecting the fact that smooth muscle fibers are part of healthy prostate tissues, whereas prostate cancer glands are per definition closely packed with less intervening stroma (Humphrey et al., 2016). This agrees with the variability observed for the DES as discussed above. Proteins associated with cell cycle-related functions such as nucleosome and chromatin assembly displayed a high degree of heterogeneity in malignant tissues. Thus, our data is consistent with recent findings suggesting that the proliferation rates among prostate cancer cells can be highly variable (Zellweger, Gunther et al., 2009), and that epigenetic events are of high importance in prostate carcinogenesis (Beharier, Shusterman et al., 2015, Grasso, Wu et al., 2012, Plass, Pfister et al., 2013).

### Spatial heterogeneity of biochemical pathways

Based on the determined protein level variance patterns described above we could also interrogate the ITH of biochemical pathways. To quantify a pathway’s variance we computed the average biological variance (intra-tissue variance) for all human pathways from ConsensusPathDB (Kamburov, Stelzl et al., 2013) with at least five quantified proteins (**Fig. 4C**). Like the individual proteins, we grouped pathways into five groups depending on their degrees of heterogeneity in malignant and benign tissues. Five pathways emerged as being particularly variable in tumor tissues (*i.e.*, average biological variance in malignant samples above 0.02): ‘Fanconi Anemia Pathway’, ‘Meiosis’, ‘Meiotic synapsis’, ‘Regulation of cell cycle progression by plk3’, as well as ‘Role of brca1 brca2 and atr in cancer susceptibility’. These pathways are involved in DNA damage response and include proteins such as serine/threonine-protein kinase ATR and the cohesion complex. The specific role of these pathways in responding to chromosomal aberrations suggests that the occurrence and repair of double strand breaks (which are a hallmark of prostate cancer) are heterogeneous within tissue specimens (Haffner, Aryee et al., 2010). Pathways highly variable only in non-tumorous tissues are markedly enriched for immune activity. The stromal component of BPH samples demonstrated a high degree of ITH in antigen processing and presentation, naïve CD8+ T cells signaling, IL12- mediated signaling, interactions between a lymphoid and a non-lymphoid cell, MHC class I complex expression, NK-cell mediated cytotoxicity, suggesting the combat between carcinogenesis and immunity. Consistent with the previous analysis, we observed more variable muscle contraction activity in non-tumorous tissues. The only pathway variable in both tumorous and non-tumorous tissues was the synthesis of phosphatidic acid, a critical component of mTOR signaling and a biosynthetic precursor for all cellular acylglycerol lipids with critical roles in prostate tissue biology (Fang, Vilella-Bach et al., 2001, Foster, 2009).

### Investigation of spatial heterogeneity of selected proteins using immunohistochemistry (IHC) in an independent cohort

We further investigated the biological variation of selected proteins from the PCT-SWATH analysis using a complementary technology in an independent, larger cohort. We constructed a tissue microarray (TMA) using benign and malignant (ADCA) prostate tissues from 83 additional patients and established IHC assays to measure the expression of ten representative proteins in the various ITH groups identified from the PCT-SWATH results, including ACTR1B, DES, PSA, GDF15 as shown in **Fig. 5**, as well as ACPP, ABCF1, NUP93, CUTA, CRAT, and FSTL1 (**Supplementary Fig. 5**). This set of validation proteins contains some well-established markers for prostate cancer in order to elucidate their variability within benign and tumorous tissue specimens. The stained TMAs contained duplicate tissue cores of 48 ADCA and 35 BPH samples. The heterogeneity of proteins was evaluated based on an immunoreactivity score computed from duplicate tissue spots and measured by the Pearson correlation coefficient between the two spots for BPH and ADCA respectively (**Fig. 5**). Thus, a high Pearson correlation score indicates a homogeneous distribution of the respective protein in the TMAs (*i.e.* low ITH). We found that the degree of ITH determined in the three patients by PCT-SWATH was well validated in the independent cohort. ACTR1B is an actin-related protein in the dynamin complex to construct cytoskeleton. This house-keeping protein exhibited a very high degree of correlation in both BPH (r = 0.96) and ADCA (r = 0.80) samples, serving as a positive control. In the TMA cohort, DES was more variable in BPH (r = 0.51) than in ADCA (r = 0.67), which is consistent with proteomics data. Our TMA data demonstrated that in BPH samples, PSA was found only in the glandular tissue, and expressed more heterogeneous than in ADCA samples, with blood PSA levels being a non-specific biomarker for PCa. Growth/differentiation factor 15 (GDF15) is a stress-induced cytokine belonging to the transforming growth factor beta superfamily (Vanhara, Hampl et al., 2012). This protein is expressed in highly complex forms with distinct biological functions related to immunity. In various tumors including prostate cancer, GDF15 interacts with the extracellular matrix and promotes tumor progression and metastasis (Vanhara et al., 2012). We found GDF15 to be expressed at relatively low levels in BPH with a low degree of ITH probably due to inflammatory changes of glandular architecture followed by stromal tissue increase in BPH (Vanhara et al., 2012). In the ADCA samples, GDF15 expression was elevated with a high degree of variation, indicating complex interactions between tumor cells and the microenviroment via modulators including GDF15. The high variability of ACPP in BPH samples was also confirmed in this cohort. Proteins grouped as medium heterogeneity including ABCF1, NUP93, CUTA, CART, and FSTL1 displayed consistent heterogeneity patterns after manual inspection of the TMA data. Taken together, we observed significant correlations between the heterogeneity measured in the TMAs and the biological variance measures obtained with PCT-SWATH across all 10 proteins (**Fig. 6**).

**Figure 5.**
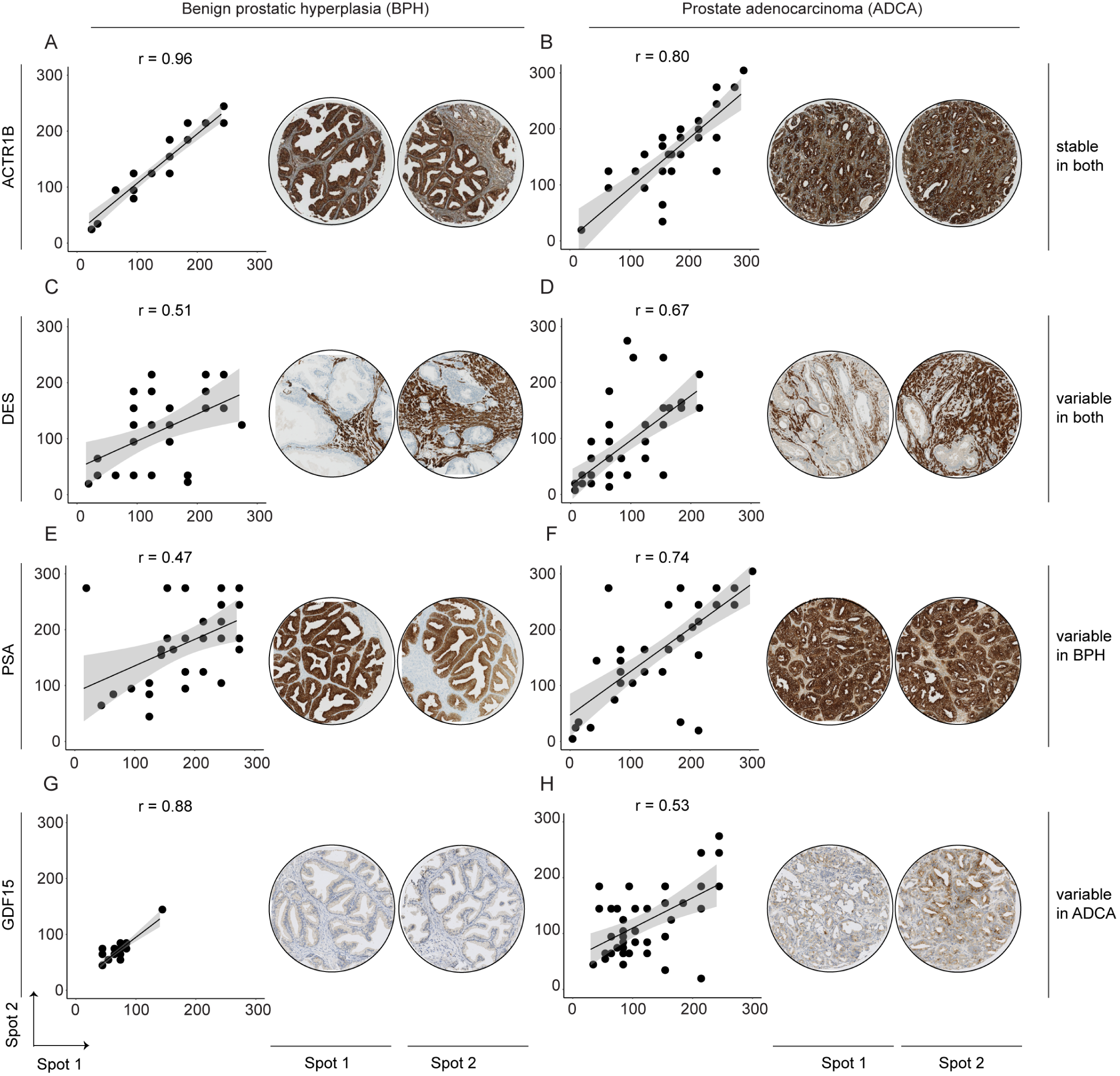
**Immunohistochemical validation of representative proteins.** The top proteins from four ITH groups in BPH and malignant (ADCA) prostate tissue were validated using a TMA with two representative tissue spots of each patient.

**Figure 6.**
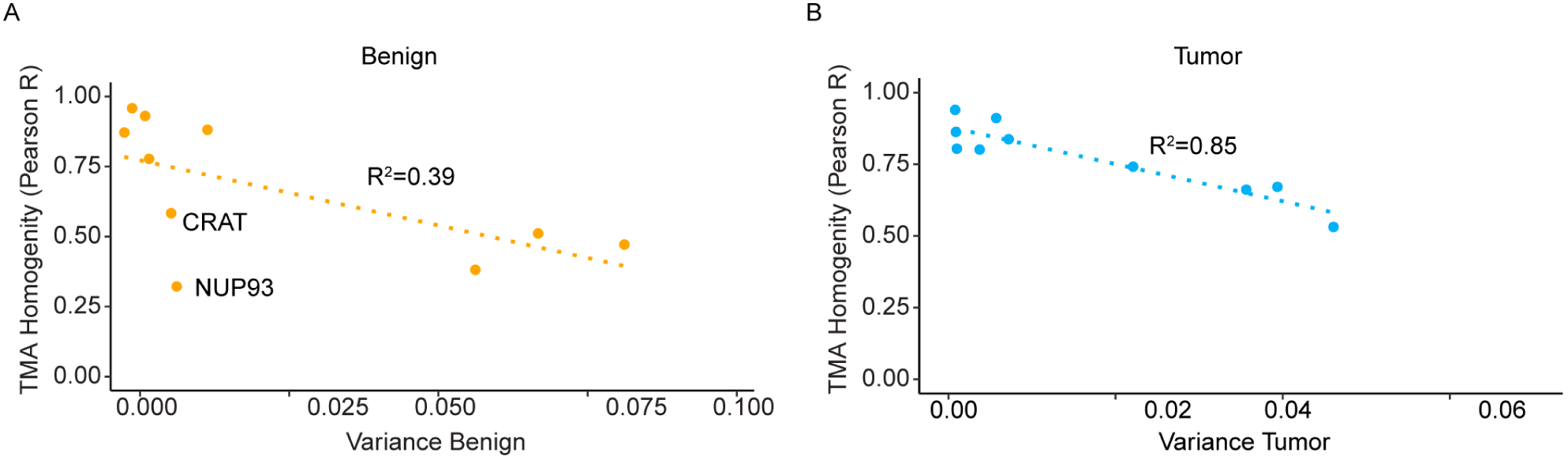
**Correlation between mass spectrometry-based (MS) variance estimates and TMA homogeneity. *A***shows benign tissues while ***B***depicts tumor tissues. The concentrations of CRAT and NUP93 were almost zero in the benign tissue samples. Thus, it is virtually impossible to estimate their intra-tissue variation in benign tissues. The correlation between MS-based variance and TMA homogeneity was however computed without excluding these two proteins. NUP93 was slightly off the regression curve because its signal in IHC was relatively weak.

## DISCUSSION

This study investigated the spatial variability of the prostate proteome, which serves as a basis for better understanding the biology of PCa protein biomarkers. Protein biomarkers including PSA and GDF15 have been well studied in PCa, however, their spatial expression in prostate tissues has not been systematically studied. ITH has been studied at the morphologic and genomic level in diverse cancers, and it poses a major challenge for cancer biology and diagnosis (Alizadeh et al., 2015). However, proteomic ITH remains underexplored in prostate cancer, despite the critical roles of proteins in tumorigenesis and cellular biochemistry in general and the various single cell-based methods.

This study represents a technical advance towards understanding spatial ITH at the proteome level for solid tumors and other tissues. Using the PCT-SWATH methodology (Guo et al., 2015b) and an associated data analysis strategy (Röst et al., 2014), we achieved deep proteomic coverage (consistent quantification of 8,248 reviewed SwissProt proteins across the 60 prostate tissue samples), and performed quantitative analysis of spatial ITH of 3,700 proteins, which were quantified by at least two proteotypic peptides that showed consistent abundance across samples. Despite the rigorous filtering, we could quantify a similar number of proteins like other studies of primary tissues, which used extensive peptide fractionation (Zhang, Wang et al., 2014, Zhang, Liu et al., 2016), and a three times higher number of proteins than a recent proteomic analysis of primary prostate tissue samples (Iglesias-Gato, Wikstrom et al., 2016). The number of proteins quantified in this study exceeds by 1-2 orders of magnitude the number of proteins typically quantified by tissue staining, which is the current standard method for protein quantification in clinical tissue samples. Our data did not achieve single cell resolution like the CyTOF technology These technologies, however, quantify orders of magnitude fewer proteins (Amir el, Davis et al., 2013, Giesen et al., 2014, Levine, Simonds et al., 2015). The data generated in this study are unique with respect to the structure of the sample set, the degree of proteomic coverage, and the degree of measurement reproducibility and accuracy. Nevertheless, new MS-based proteomics technologies enabling analysis of single cells from tissue samples will be desirable to quantify spatial ITH at higher spatial resolution in future studies.

The main goal of this study was not to discover new protein biomarkers; instead we aimed to characterize the spatial ITH of the prostate proteome and investigate whether the ITH influences the utility of protein biomarkers and candidates. Our data contributed to the understanding of the following prostate cancer biology. First, we systematically reported the degree of ITH of 3,700 proteins in prostate tissues. Although some of these proteins are widely used in clinic, their expression pattern in prostate tumors was unclear. We found PSA preferentially variable in BPH, while GDF15 tended to vary in different tumor regions. This finding, together with the ITH pattern of eight more clinically relevant protein biomarkers, were further investigated and confirmed in an independent cohort of 83 PCa patients using TMA technology. This additional cohort analysis not only confirm that the PCT-SWATH technology is a valid and practical extension of IHC and TMA for proteome-scale ITH analysis of clinical tissue samples, but also consolidated the spatial variability of these proteins in prostate tissues, providing guidance for clinical application of these proteins as biomarkers. We found protein ITH patterns to vary between tissue types due to their biological functions and interplay with the microenvironment.

Second, the data also shed light on the heterogeneity of multiple biochemical pathways. Interestingly, benign tissue displayed a high degree of variability in immunity-related signaling pathways, whereas tumor tissues, characterized by enhanced proliferation and DNA-damage, exhibited high degree of heterogeneity in several DNA damage response pathways, suggesting that spatially variable DNA repair pathways probably contributed to genomic heterogeneity during the evolution of prostate cancers. Further, we found that the degree of intra-tissue variability of multiple pathways was slightly higher in benign specimens compared to malignant tissues (**Fig. 4**), which may be due to the more complex structure of healthy tissues involving a larger number of distinct cell types, while in tumorous tissues most cell types are replaced by tumor cells.

The observed intra-tissue protein variability patterns have implications that extend beyond the present study to protein biomarker studies in general and have specific significance for biomarker studies in the context of personalized medicine, where sample availability is generally sparse. Our data suggest that the variation of some protein levels between patients is similar in magnitude to the variation within a single prostate. These findings underline the significance of low intra-tissue variability as an important property of a clinical protein biomarker. In fact, the observed variability patterns provide a rational explanation why some previously published tissue biomarker studies did not produce concordant results. Similar conclusions were drawn in an earlier study, in which the abundance variability of plasma proteins was analyzed in a twin cohort (Liu, Buil et al., 2015). The data indicated that those biomarker candidates that were proposed in the literature and eventually approved for clinical use showed low levels of variability derived from genetic differences in a population. In contrast, biomarker candidates proposed in the literature that showed a high degree of genetically caused abundance variation in a population were rarely validated. Our data add a new perspective to this problem: a candidate biomarker may show high variability between patients when quantified using single needle biopsies per patient. However, the tumor-wide average concentrations may not be substantially different, and the true cause of the apparent inter-patient variability may be ITH, rather than rooted in the biochemical difference between normal and tumor tissues. Therefore, we suggest that intra-tissue variability of a protein or a pathway be used as an important criterion for the assessment of protein biomarker candidates, in addition to other parameters such as expression level and biochemical function. Including more biological replicates per patient to average out protein ITH or increasing patient numbers to account for variability may not always be possible. Thus, our work provides an important lead as to how ITH can be tackled even for small patient and sample numbers in clinically realistic scenarios.

## MATERIALS & METHODS

### Patients and samples for PCT-SWATH analyses

The prostates from three patients after prostatectomy were cut into tissue sections (thickness: about 3 mm). Fresh BPH and ADCA tissue sections were frozen and embedded in O.C.T.. The tissue were examined by trained pathologists and graded similarly according to the Gleason system as shown in **Fig. 1**. Tumorous tissues from each patient contained acinar prostate tumors, while one patient included an extra ductal prostate tumor. To obtain biopsy-scale tissue samples for PCT-SWTH analysis, we utilized a needle to punch out tissue cylinders (diameter: 1 mm, length: ∼ 3 mm) at the locations as shown in **Fig. 1**. Multiple (three or six) punches were obtained from each area. The Ethics Committee of the Canton of Zurich approved all procedures involving human fresh frozen material. All three patients were part of the Zurich prostate cancer outcomes cohort study (ProCOC, KEK-ZH-No. 2008-0040) (Umbehr, Kessler et al., 2008, Wettstein, Saba et al., 2017), and each patient signed an informed consent form.

### PCT-SWATH

The tissue samples were first washed to eliminate O.C.T., followed by PCT-assisted tissue lysis and protein digestion, and SWATH-MS analysis, as described previously (Guo et al., 2015b). Briefly, each tissue punch was washed with 70% ethanol / 30% water (30 s), water (30 s), 70% ethanol / 30% water (5 min, twice), 85% ethanol / 15% water (5 min, twice), and 100% ethanol (5 min, twice). Subsequently, the tissue punches were placed in PCT-MicroTubes with PCT-MicroPestle and 30 μl lysis buffer containing 8 M urea, 0.1 M ammonium bicarbonate, Complete protease inhibitor cocktail (Roche) using a barocycler (model NEP2320-45k, PressureBioSciences, South Easton, MA) which offers cycling alternation of high pressure (45,000 p.s.i., 50 s per cycle) and ambient pressure (14.7 p.s.i., 10 s per cycle) for 1 h. The extracted proteins were then reduced and alkylated prior to lys-C and trypsin-mediated proteolysis under pressure cycling. Lys-C (Wako; enzyme-to-substrate ratio, 1:40) ‐mediated proteolysis was performed under 45 cycles of pressure alternation (20,000 p.s.i. for 50 s per cycle and 14.7 p.s.i. for 10 s per cycle), followed by trypsin (Promega; enzyme-to-substrate ratio, 1:20)-mediated proteolysis using the same cycling scheme for 90 cycles. The resultant peptides were cleaned by SEP-PAC C18 (Waters Corp., Milford, MA) and analyzed, after spike-in 10% iRT peptides, using SWATH-MS following the 32-fixed-size-window scheme as described previously with a 5600 TripleTOF mass spectrometer (Sciex) and a 1D+ Nano LC system (Eksigent, Dublin, CA). The LC gradient was formulated with buffer A (2% acetonitrile and 0.1% formic acid in HPLC water) and buffer B (2% water and 0.1% formic acid in acetonitrile) through an analytical column (75 μm × 20 cm) and a fused silica PicoTip emitter (New Objective, Woburn, MA, USA) with 3-μm 200 Å Magic C18 AQ resin (Michrom BioResources, Auburn, CA, USA). Peptide samples were separated with a linear gradient of 2% to 35% buffer B over 120 min at a flow rate of 0.3 μl min^1^. Ion accumulation time for MS1 and MS2 was set at 100 ms, leading to a total cycle time of 3.3 s.

### SWATH assays for prostate tissue proteome

We also analyzed unfractionated prostate tissue digests prepared by the PCT method using Data Dependent Acquisition (DDA) mode in a tripleTOF mass spectrometer over a gradient of 2 hours as described previously (Röst et al., 2014). We spiked iRT peptides (Escher, Reiter et al., 2012) into each sample to enable retention time calibration among different samples. We then combined this library with the DDA files from pan-human library (Rosenberger, Koh et al., 2014). All together we analyzed 422 DDA files using X!Tandem (MacLean, Eng et al., 2006) and OMSSA (Geer, Markey et al., 2004) against a target-decoy, non-redundant human UniProtKB/Swiss-Prot protein database (Oct 21, 2016) containing 20,160 protein sequences and the iRT peptide sequences. Reversed protein sequences were used as decoy sequences. We allowed maximal two missed cleavages for fully tryptic peptides, and 50 p.p.m. for peptide precursor mass error, and 0.1 Da for peptide fragment mass error. Static modification included carbamidomethyl at cysteine, while variable modification included oxidation at methionine. Search results from X!Tandem and OMSSA were further analyzed through Trans-Proteomic Pipeline (TPP, version 4.6.0) (Deutsch, Mendoza et al., 2010) using PeptideProphet and iProphet, followed by SWATH assay library building procedures as detailed previously (Guo et al., 2015b, Schubert, Gillet et al., 2015). Altogether, we identified 160,442 peptides with <1% FDR.

### Peptide quantification using OpenSWATH

SWATH files were analyzed using the prostate tissue proteome assay library described above and OpenSWATH software as described previously (Röst et al., 2014). Briefly, wiff files were converted into mzXML files using ProteoWizard msconvert v.3.0.3316, and then mzML files using OpenMS (Sturm, Bertsch et al., 2008) tool FileConverter. OpenSWATH was performed using the tool OpenSWATHWorkflow with input files including the mzXML file, the TraML library file, and TraML file for iRT peptides. The false discovery rate for peptide identification was below 0.1%. High confidence peptide features from different samples were aligned using the algorithm TRansition of Identification Confidence (TRIC) (version r238), which is available from https://pypi.python.org/pypi/msproteomicstools or https://code.google.com/p/msproteomicstools. The following parameters for the feature_alignment.py are as follows: maxrtdiff = 30, method = global_best_overall, nr_high_conf_exp = 2, target_fdr = 0.001, use_score_filter = 1.

### Protein quantification

The concentration of each protein was quantified through the simultaneous measurement of several peptides. To optimize the protein quantification, we developed a new computational method, which combines maximally consistent peptides for each protein and excludes inconsistent (*i.e.* uncorrelated) peptides (Picotti et al., 2013). For example, variation of post-translational modifications (PTM) would result in peptide level variation that is uncorrelated across samples, because mostly only one of the two peptides would be affected by the PTM.(Picotti et al., 2013). Given a set of peptides unambiguously assigned to a single protein, consistent peptides were selected using the following procedure: all pairwise correlations between all peptides of a protein across the samples were calculated at first. Peptide pairs with a Pearson correlation coefficient (R) of at least 0.3 were determined, resulting in clusters of correlated peptides. This procedure yielded one or more peptide clusters per protein. We used the largest cluster of each protein and we quantified the protein’s concentration as the average intensity across the peptides in that cluster. The minimum cluster size was set to 2 and proteins without a cluster of at least two correlated peptides were removed from the subsequent analysis. This procedure resulted in very robust concentration estimates for 3,700 proteins with high correlation between technical replicates (R ≥ 0.95) and no missing values.

### Determining the biological variance between punches in a specific tissue (intra-tissue variance)

Measurements of protein abundance differences between individual punches are affected by a combination of biological and technical factors. Thus, to quantify the biological variation between punches we need to subtract the technical variance from the total variance, *i.e.* the combined variance due to technical and biological factors. Estimating the biological variance of protein levels between punches therefore requires estimates of the technical variance and the total variance. Intuitively, one would estimate both variances using a standard approach such as ANOVA in a single statistical model. However, technical replicates are paired because they come from the same punch and thus they are not independent, whereas the total variance needs to be estimated across punches, *i.e.* involving partially independent measurements.

Therefore we decided to separately estimate technical and total variances. Here, technical variance was estimated from the dispersion of measurements between paired technical replicates and total variance was estimated from the dispersion of measurements between independent punches from the same specimen. Compared to an approach estimating both technical and total variance in a single statistical model, our approach has the caveat that the two variance estimates can be inconsistent in the sense that the estimated total variance can be smaller than the estimated technical variance. Obviously, this happens only for those proteins where the technical noise is large compared to the biological variance, in which case it is anyways impossible to reliably estimate the true biological variance (no matter which statistical approach is taken). We therefore conservatively accept that in those cases we cannot provide an estimate of the biological variance. However, we assume that in most of those cases the biological variance will be small compared to the other proteins for which we could estimate a biological variance.

In detail, the variances were estimated in the following way.

First, the protein concentrations (computed from peptide intensities as described above) were log10-transformed. Next, protein concentrations were quantile normalized per sample. As the signal distributions between non-tumorous (benign) and tumorous tissue (malignant: acinar and ductal) differed significantly, the normalization was performed separately for each tissue type. For each protein, we computed the technical variation for each sample and averaged the inter-replicate variance across all 30 samples (Tukey, 1977). Since technical replicates are (obviously) paired, the technical variance was estimated as the dispersion of the two replicates from their sample mean averaged across all punches *(n =* 30). Thus, the technical variance *VAR_TECH_* of protein *i* was estimated as:

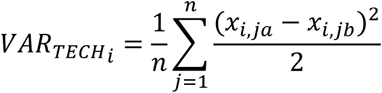

with *x_i,ja_* and *x_i,jb_* being the two technical replicates (*a* and *b*) of the protein level measurements from punch *j*. In this case, no batch correction was performed, because batch correction would reduce the technical variance (technical replicates were always in different batches), which might lead to underestimation of the technical variance. The final estimate of technical variances was computed after removing outliers above and below the 1.5*IQR of 30 samples based on Tukey’s method (Tukey, 1977).

The total variances between punches (*i.e.* the combined variance from technical noise and biological variance) were initially computed for each batch separately. Thus, variation among punches from the same specimen (same patient *p* and same tissue type *t*) were averaged. Finally, total variances *VAR_TOT_* between punches were averaged across batches.

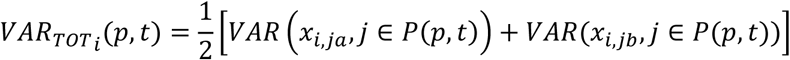

Where *P*(*p,* t) denotes all punches *j* from patient *p* and tissue *t* (i.e. either benign, acinar, or ductal). The indices *a* and *b* denote the two technical replicates, as above. Thus, total variances were estimated purely from deviations *within* batches and are (unlike technical variances) not affected by batch-to-batch variation. As a consequence, technical variances are biased towards larger values compared to total variances. This approach is conservative in the sense that it minimizes the number of proteins that are falsely classified as having variable concentrations within tissues. Thus, this approach will likely underestimate the true number of proteins with large biological intra-tissue variance. Given the total variance and technical variance, the biological variance *VAR_BIO_* of protein *i* was computed as follows:

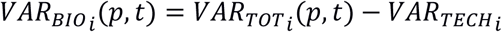

This scheme generated seven independent estimates of total variance per protein: four for the patients no. 1 and no. 2 (benign and malignant acinar tissues) and three for patient no. 3 (benign, acinar, and ductal). The intra-tissue variance shown in Figure 4 is the average biological variance of a given protein across all patients and tissue types. The tissue-specific variances used for Figure 5 are the average variances across the patients for the respective tissues (benign, acinar, ductal). The biological variance in tumor was estimated as the average of all acinar and the ductal (patient 3) tumor regions.

### Grouping of proteins and pathways based on their variability

In cases where the estimated technical variance is greater than the estimated total variance, subtracting the technical from the total variance yields a negative ‘variance estimate’ (**Supplementary Fig 4**). Because these negative ‘variances’ are the result of our imperfect variance estimates, the distribution of these values can be used to quantify the inherent uncertainty in our estimates of the biological variance. Thus, we can use the distribution of the absolute values (the ‘mirror distribution’ into the positive range) as a background distribution for the Null hypothesis that the true biological variance is indistinguishable from zero (*or:* that the total observed variance is exclusively due to technical variance). Based on this approach, 797 proteins had p-values below 0.01 and were thus classified as biologically variable proteins (*i.e.* significantly variable within the same specimen). These797 variable proteins were further sub-classified as follows: if the ratio *of biological variance in benign* to *biological variance in tumor* was above 2 they were classified as “variable in non-tumor” (339 proteins); if the ratio of *biological variance in tumor* to *biological variance in normal* was above 2, proteins were classified as “variable in tumor” (93 proteins); 365 proteins with similar variances in both tissue types *(i.e.* not different by more than a factor of 2) were classified as “variable in non-tumor and tumor*”.* Stable proteins were defined by choosing the 100 proteins with the lowest biological variance. Remaining proteins, which were not assigned to any of the above four groups, were classified as “medium heterogeneity” proteins.

Note that our computation of empirical p-values for determining variable proteins is not critical for the conclusions. If we had simply chosen the top 200 most variable proteins (as the basis for groups 1-3) and compared them to the 200 most stable proteins (group 4) the conclusions would be virtually identical.

### Gene Set Enrichment Analysis

Gene Ontology (GO) enrichment of proteins was performed using topGO, which takes the topology of the ontology into account. The enrichment analysis was carried out by using Fisher’s exact test with the background of measured proteins in this study. We excluded GO terms with less than 10 proteins and with more than 300 proteins from the analysis (the former are too small, the latter are too generic). Further, we reported only GO terms that had at least 4 proteins enriched (overlapping).

Intra-tissue heterogeneity of entire biochemical pathways was determined according to the protein level variance. Pathway variability was calculated by averaging the biological variances of all proteins annotated for a given ConsensusPathDB pathway. We required that each pathway contained at least five quantified proteins. ConsensusPathDB combines pathway annotations from different sources. Thus, in some cases the same pathway is reported more than ones. In such case the pathway variant with the largest number of quantified proteins was used.

### Determining the variance between tissues (inter-tissue variance) and between patients (inter-patient variance)

Batch effects were corrected by centering each protein’s concentration per batch. In our experimental design, batches were balanced in the sense that each batch had the same number of benign and malignant samples (3 of each) and each batch had the same number of samples from the same patient (2 patients per batch, 3 samples from each patient).

Inter-tissue variances were estimated using concentrations centered per patient (subtracting patient mean). Inter-patient variances were estimated using concentrations centered per tissue type (subtracting tissue mean across patients). All of those computations were based on batch-corrected concentrations and after averaging technical replicates. Batch-corrected values were also used for Figure 2.

### Patient cohort and tissue microarray (TMA)

The Ethics Committee of the Kanton St. Gallen, Switzerland approved all procedures involving human materials used in this TMA, and each patient signed an informed consent. For the study, patients with BPH and matching ADCA were included, whereas advanced prostate cancer, infectious or inflammatory diseases, or other malignancies fulfilled exclusion criteria as described previously (Cima, Schiess et al., 2011). A TMA was constructed using formalin-fixed, paraffin-embedded tissue samples derived from 83 patients (BPH, n = 35; ADCA, n = 48).

### Immunohistochemical staining and evaluation

The following primary antibodies were used to stain 4 μm slides of the TMA using the Ventana Benchmark (Roche Ventana Medical Systems, Inc., Tucson, AZ, USA) automated staining system: ACTR1B (1:400; Abcam, 60 min pretreatment), Desmin/DES (1:20; DAKO A/S, 16 min pretreatment), KLK3/PSA (1: 10000; DAKO A/S) and GDF15 (1:50; biorbyt, 30 min pretreatment), ACPP (1:2000; DAKO A/S), ABCF1 (1:50; Novus Biologicals, 90 min pretreatment), NUP93 (1:50; NovusBiologicals, 60 min pretreatment), CUTA (1:100; LifespanBiosciences, 60 min pretreatment), CRAT (1:100; Atlas Antibodies, 30 min pretreatment), and FSTL1 (1:100; Atlas Antibodies, 16 min pretreatment). Detection was performed with ChromoMap Kit (Ventana) for ABCF1, PCP4, CUTA and OptiView DAB Kit (Ventana) for the others (Desmin, KLK3/PSA, NUP93, CRAT, FSTL1,PAP) using the heat-induced epitope retrieval CC1 solution. Slides were counterstained with hematoxylin (Ventana), dehydrated and mounted. For GDF15 4 μm slides were stained using the Leica Bond (Leica Biosystems, Muttenz, Switzerland) automated staining system. For detection the Bond Polymer Refine Detection kit and heat-induced epitope retrieval HIER2 solution (Leica Biosystems) following Hematoxylin counterstaining was used. Staining intensities for each antibody were evaluated in a semi-quantitative, 4-tier manner (negative = 0, weak = 1, moderate = 2 and strong = 3), along with the occupied area (in 1%, 3%, 5% and above 10% steps), by one pathologist (N.J.R.). An immunoreactivity score (IRS; staining intensity multiplied by percentage of spot; similar to the recommendations by Remmele & Stegner (Remmele & Stegner, 1987) consisting of “staining intensity x area (%)” was calculated.

### Data deposition

The SWATH raw data and analyzed data as well as assay library are deposited in PRIDE (Vizcaino et al., 2014). For the SWATH data of the three patients: Project accession: PXD003497; Username; Password: Vvl6EFPj. For the SWATH data of the 27 patients: Project accession: PXD004589; Username;Password: 1zHGceA9.

## Acknowledgements

This work was supported by the SystemsX.ch project PhosphoNet PPM (to R.A. and P.J.W.), the Swiss National Science Foundation (grant no. 3100A0-688 107679 to R.A.), the Foundation for Research in Science and the Humanities at the University of Zurich (to P.J.W.), the European Research Council (grant no. ERC-2008-AdG 233226 and ERC-2014-AdG670821 to R.A.), and the German Federal Ministry of Education and Research (BMBF; grants: Sybacol & PhosphoNetPPM to L.L. and A.B.). We thank O.L. Kon for critical reading of the manuscript.

## Author contributions

A.B., T.G., P.J.W. and R.A. designed the project. P.J.W., Q.Z., C.E.W., C.P, C.F, K.S., J.H.R. and N.J.R. procured the samples. T.G. performed the PCT-SWATH analysis. L.L., K.C. and A.B. designed and performed the statistical analyses. L.L., A.B., T.G. and R.A. interpreted the results. P.J.W., C.F., N.J.R., Q.Z., U.W. and W.J. performed tissue microarray validation. T.G., L.L., A.B., Q.Z., and R.A. wrote the manuscript with inputs from all co-authors. R.A., A.B. and P.J.W. supported and supervised the project.

## SUPPLEMENTARY FIGURES

**Supplemental Figure 1.**
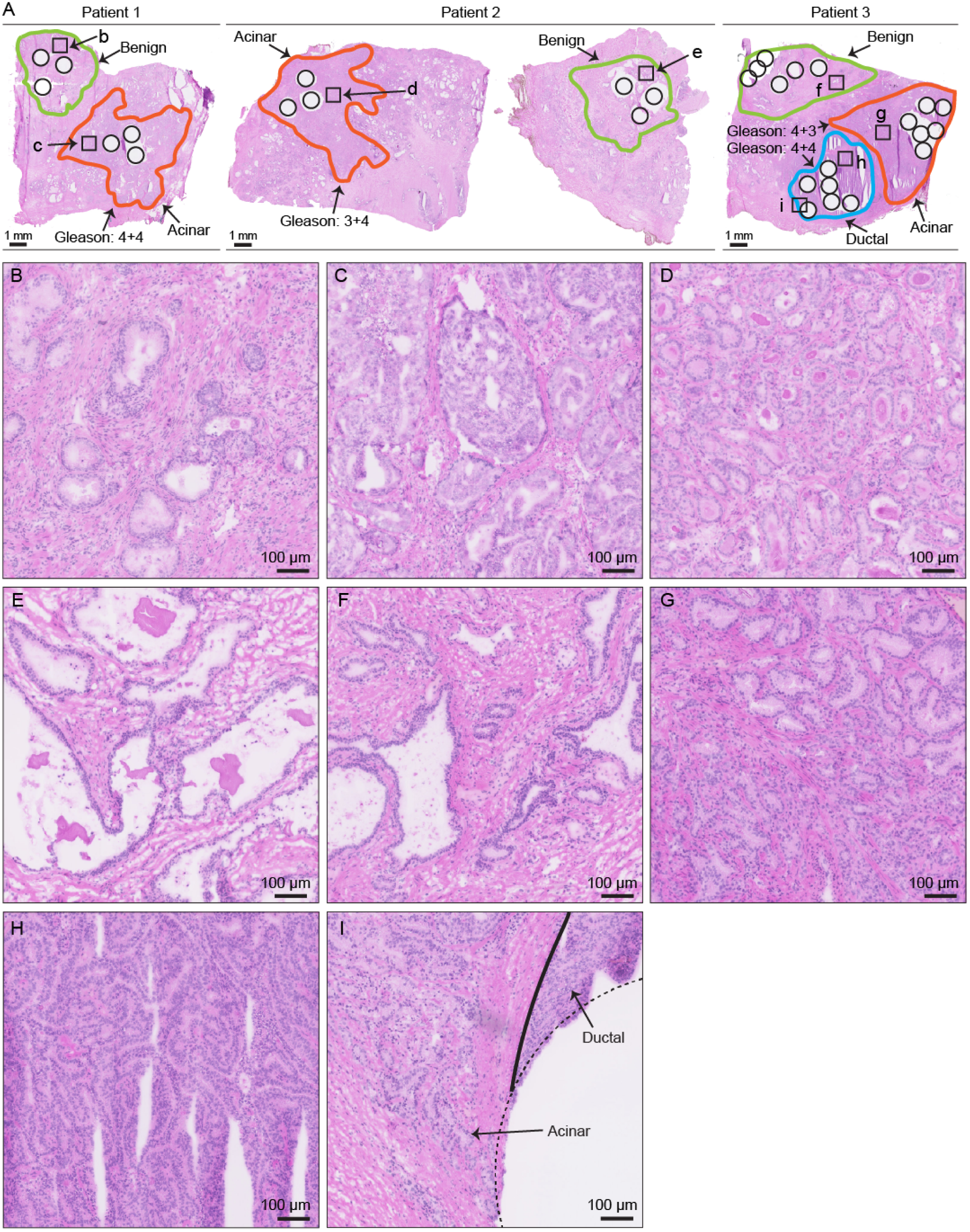
**Benign and malignant prostate tissue from three individuals.** H&E staining of the fresh frozen prostate tissue used in this study. Amplified views of representative region in each area were shown in **B** – **I** as indicated.

**Supplemental Figure 2.**
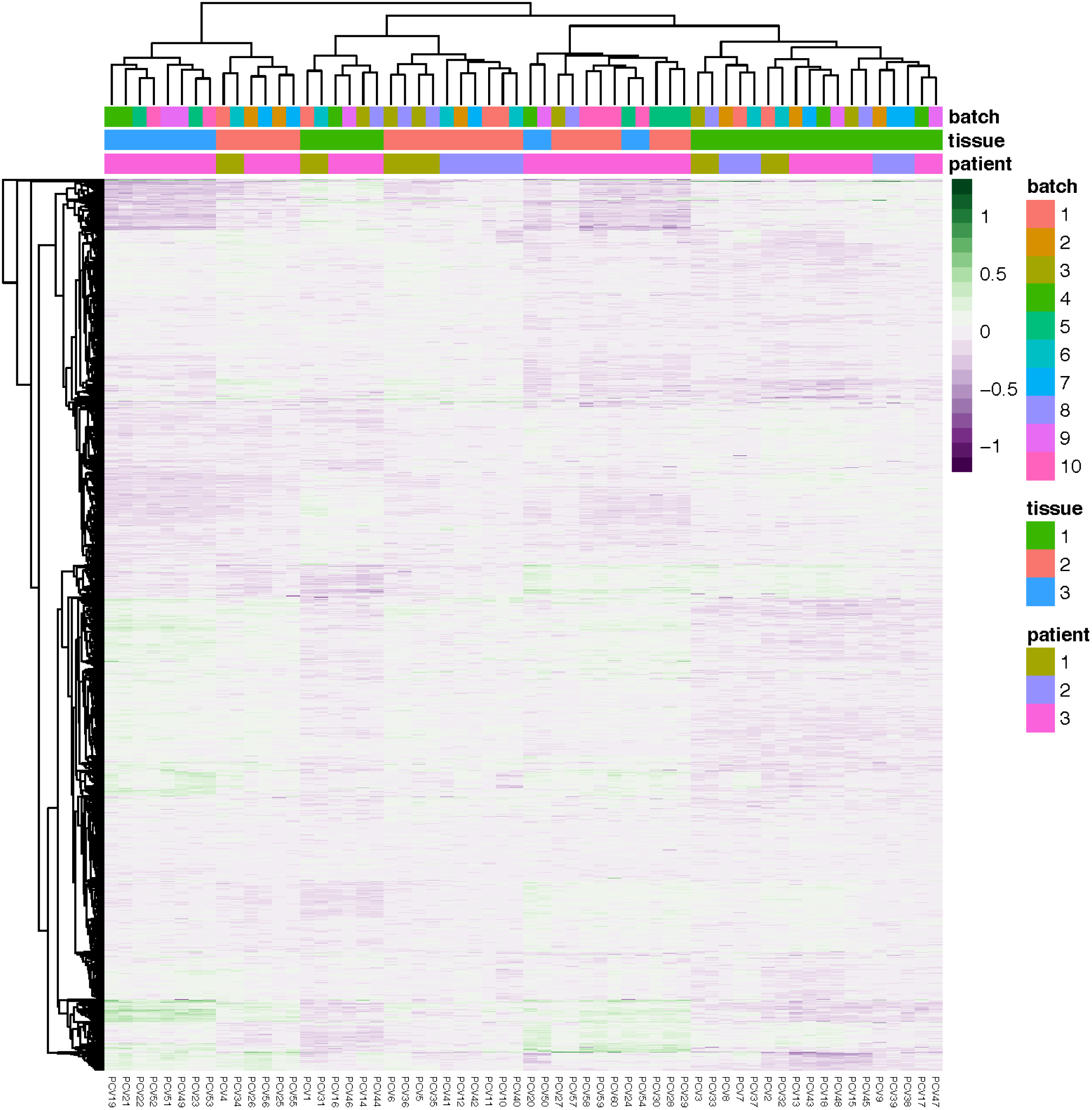
**Unsupervised clustering of 3700 proteins quantified with at least two concordant peptides.**

**Supplemental Figure 3.**
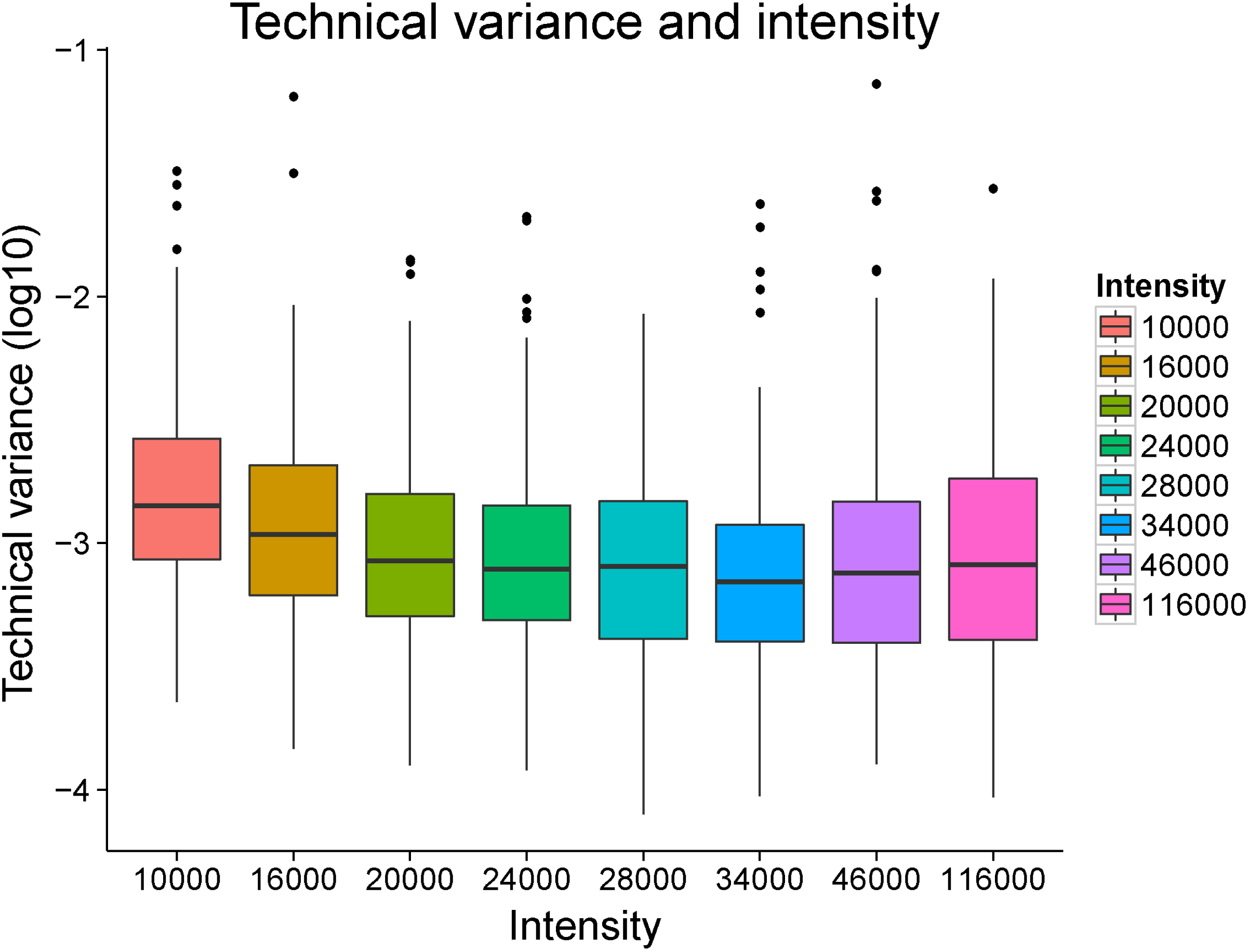
**Dependence of technical variance on protein intensity.** Proteins are divided into eight bins with roughly the same number. X-axis shows the mean intensity value of each bin, and Y-axis shows the log10 technical variance.

**Supplemental Figure 4.**
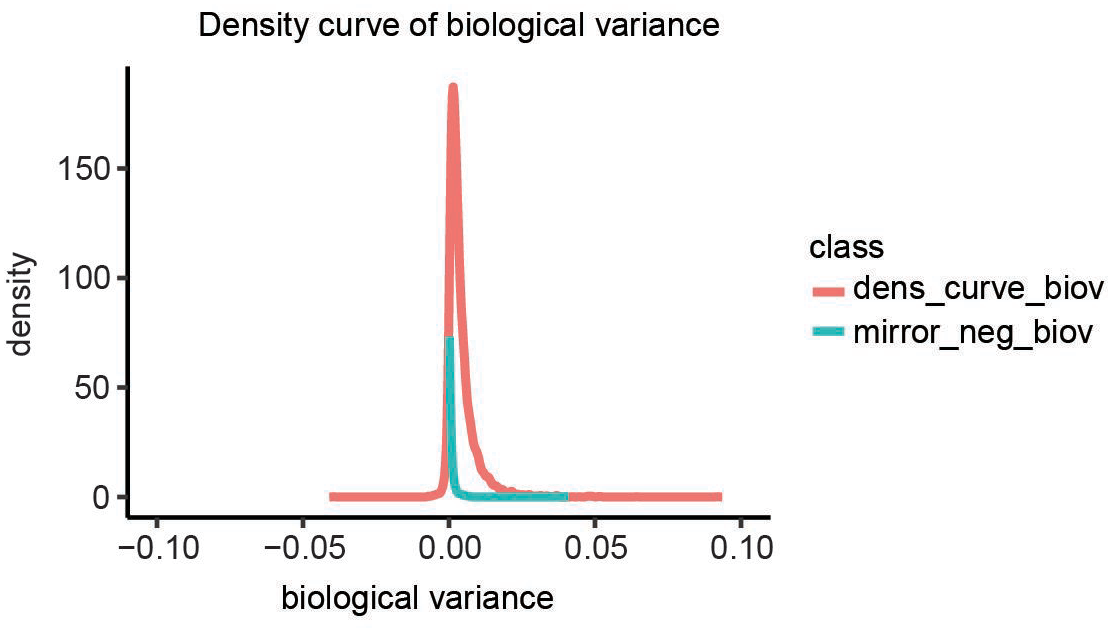
**Density curves of biological variance.** Occasionally, our estimate of the technical variance was larger than the variation between punches, after technical replicates were averaged per punch. This resulted in a negative estimate of the biological variance, which is of course infeasible. We assumed that those proteins have a biological variance close to zero, thus the total variance is mostly reflecting technical variance. Therefore, we used the distribution of negative scores as a background distribution (Null distribution) for the Null hypothesis that there is no biological variance between punches. The blue curve shows the negative part of the distribution mirrored on the positive side. The distribution of observed biological variance estimates (red) is clearly above that background distribution.

**Supplemental Figure 5.**
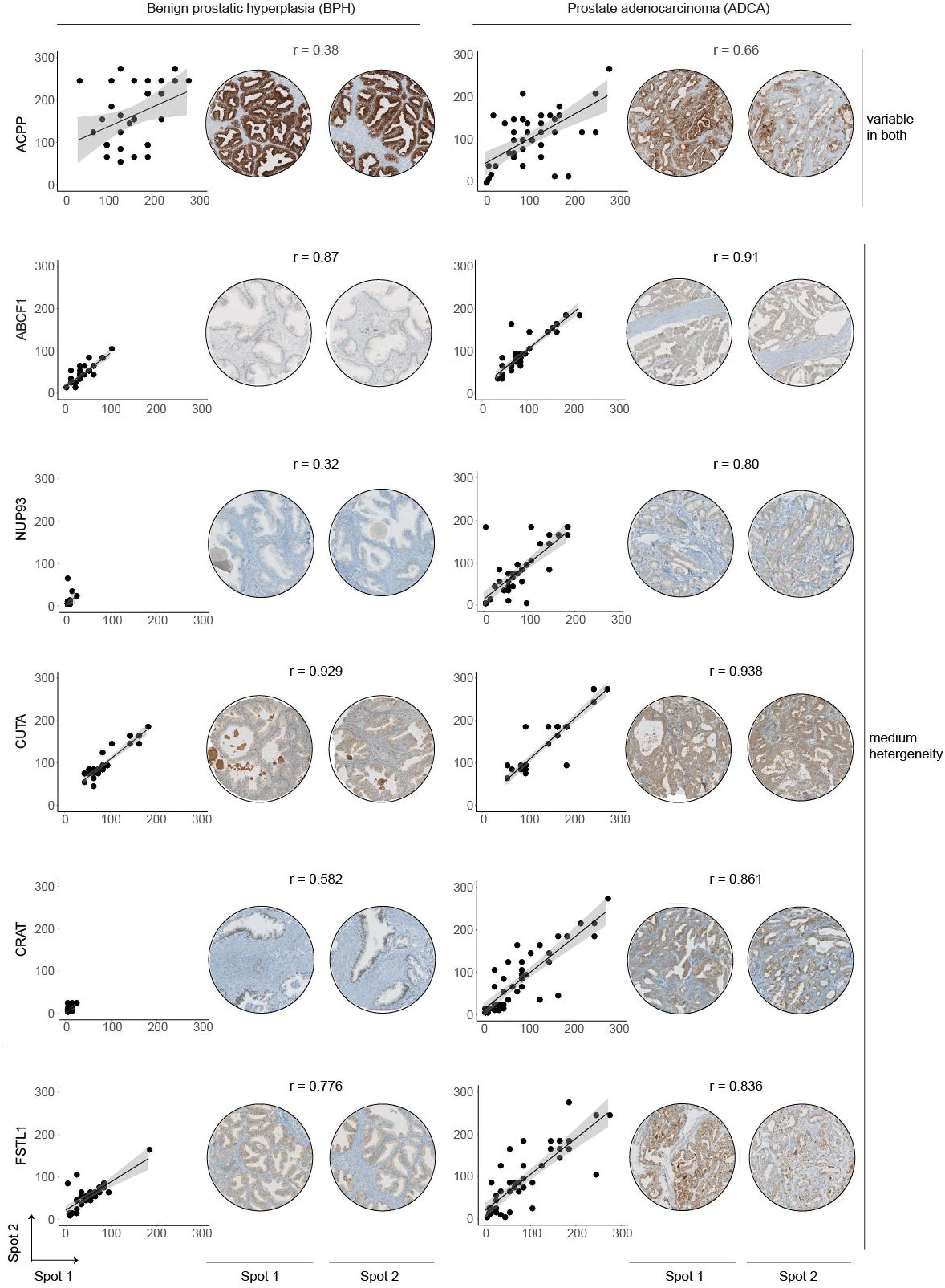
**Staining images of protein expression for the tissue microarray.** Six proteins (ACPP; ABCF1; NUP93; CUTA; CRAT; FSTL1) were measured using immunohistochemistry in a TMA containing tissue samples from 83 patients from an independent cohort, including 35 patients with BPH and 48 patients with prostate ADCA.

**Supplementary Table 1.** Annotations of the 60 prostate tissue samples used for the PCT-SWATH analysis.

**Supplementary Table 2.** Batch design of the 60 prostate tissue samples in the PCT-SWATH analysis.

**Supplementary Table 3.** Peptides identified in the 60 prostate tissue samples.

**Supplementary Table 4.** Proteins quantified in the 60 prostate tissue samples.

**Supplementary Table 5.** Biological variance of proteins in the 60 prostate tissue samples.

## References

Alizadeh AA, Aranda V, Bardelli A, Blanpain C, Bock C, Borowski C, Caldas C, Califano A, Doherty M, Elsner M, Esteller M, Fitzgerald R, Korbel JO, Lichter P, Mason CE, Navin N, Pe’er D, Polyak K, Roberts CW, Siu L et al. (2015) Toward understanding and exploiting tumor heterogeneity. Nat Med 21: 846–53.

Amir el AD, Davis KL, Tadmor MD, Simonds EF, Levine JH, Bendall SC, Shenfeld DK, Krishnaswamy S, Nolan GP, Pe’er D (2013) viSNE enables visualization of high dimensional single-cell data and reveals phenotypic heterogeneity of leukemia. Nat Biotechnol 31: 545–52.

Ayala G, Tuxhorn JA, Wheeler TM, Frolov A, Scardino PT, Ohori M, Wheeler M, Spitler J, Rowley DR (2003) Reactive stroma as a predictor of biochemical-free recurrence in prostate cancer. Clin Cancer Res 9: 4792–801.

Balk SP, Ko YJ, Bubley GJ (2003) Biology of prostate-specific antigen. J Clin Oncol 21: 383–91.

Barry MJ (2009) Screening for prostate cancer--the controversy that refuses to die. The New England journal of medicine 360: 1351–4.

Beharier O, Shusterman E, Szaingurten-Solodkin I, Weintraub AY, Sheiner E, Swissa SS, Gitler D, Hershkovitz R (2015) Placental growth factor concentration in maternal circulation decreases after fetal death: lessons from a case series study. Archives of gynecology and obstetrics 292: 1027–32.

Boutros PC, Fraser M, Harding NJ, de Borja R, Trudel D, Lalonde E, Meng A, Hennings-Yeomans PH, McPherson A, Sabelnykova VY, Zia A, Fox NS, Livingstone J, Shiah YJ, Wang J, Beck TA, Have CL, Chong T, Sam M, Johns J et al. (2015) Spatial genomic heterogeneity within localized, multifocal prostate cancer. Nat Genet 47: 736–45.

Cancer Genome Atlas Research N (2013) Genomic and epigenomic landscapes of adult de novo acute myeloid leukemia. The New England journal of medicine 368: 2059–74.

Cima I, Schiess R, Wild P, Kaelin M, Schuffler P, Lange V, Picotti P, Ossola R, Templeton A, Schubert O, Fuchs T, Leippold T, Wyler S, Zehetner J, Jochum W, Buhmann J, Cerny T, Moch H, Gillessen S, Aebersold R et al. (2011) Cancer genetics-guided discovery of serum biomarker signatures for diagnosis and prognosis of prostate cancer. Proc Natl Acad Sci U S A 108: 3342–7.

Cyll K, Ersvaer E, Vlatkovic L, Pradhan M, Kildal W, Avranden Kjaer M, Kleppe A, Hveem TS, Carlsen B, Gill S, Loffeler S, Haug ES, Waehre H, Sooriakumaran P, Danielsen HE (2017) Tumour heterogeneity poses a significant challenge to cancer biomarker research. Br J Cancer 117: 367–375.

Dalerba P, Cho RW, Clarke MF (2007) Cancer stem cells: models and concepts. Annu Rev Med 58: 267–84.

Deutsch EW, Mendoza L, Shteynberg D, Farrah T, Lam H, Tasman N, Sun Z, Nilsson E, Pratt B, Prazen B, Eng JK, Martin DB, Nesvizhskii AI, Aebersold R (2010) A guided tour of the Trans-Proteomic Pipeline. Proteomics 10: 1150–9.

Di Lorenzo G, Buonerba C, Kantoff PW (2011) Immunotherapy for the treatment of prostate cancer. Nat Rev Clin Oncol 8: 551–61.

Ding L, Ley TJ, Larson DE, Miller CA, Koboldt DC, Welch JS, Ritchey JK, Young MA, Lamprecht T, McLellan MD, McMichael JF, Wallis JW, Lu C, Shen D, Harris CC, Dooling DJ, Fulton RS, Fulton LL, Chen K, Schmidt H et al. (2012) Clonal evolution in relapsed acute myeloid leukaemia revealed by whole-genome sequencing. Nature 481: 506–10.

Domon B, Aebersold R (2010) Options and considerations when selecting a quantitative proteomics strategy. Nat Biotechnol 28: 710–21.

Epstein JI, Egevad L, Amin MB, Delahunt B, Srigley JR, Humphrey PA, Grading C (2016) The 2014 International Society of Urological Pathology (ISUP) Consensus Conference on Gleason Grading of Prostatic Carcinoma: Definition of Grading Patterns and Proposal for a New Grading System. The American journal of surgical pathology 40: 244–52.

Escher C, Reiter L, MacLean B, Ossola R, Herzog F, Chilton J, MacCoss MJ, Rinner O (2012) Using iRT, a normalized retention time for more targeted measurement of peptides. Proteomics 12: 1111–21.

Fang Y, Vilella-Bach M, Bachmann R, Flanigan A, Chen J (2001) Phosphatidic acid-mediated mitogenic activation of mTOR signaling. Science 294: 1942–5.

Foster DA (2009) Phosphatidic acid signaling to mTOR: signals for the survival of human cancer cells. Biochim Biophys Acta 1791: 949–55.

Geer LY, Markey SP, Kowalak JA, Wagner L, Xu M, Maynard DM, Yang X, Shi W, Bryant SH (2004) Open mass spectrometry search algorithm. J Proteome Res 3: 958–64.

Gerlinger M, Rowan AJ, Horswell S, Larkin J, Endesfelder D, Gronroos E, Martinez P, Matthews N, Stewart A, Tarpey P, Varela I, Phillimore B, Begum S, McDonald NQ, Butler A, Jones D, Raine K, Latimer C, Santos CR, Nohadani M et al. (2012) Intratumor heterogeneity and branched evolution revealed by multiregion sequencing. The New England journal of medicine 366: 883–92.

Giesen C, Wang HA, Schapiro D, Zivanovic N, Jacobs A, Hattendorf B, Schuffler PJ, Grolimund D, Buhmann JM, Brandt S, Varga Z, Wild PJ, Gunther D, Bodenmiller B (2014) Highly multiplexed imaging of tumor tissues with subcellular resolution by mass cytometry. Nat Methods 11: 417–22.

Gillet LC, Navarro P, Tate S, Rost H, Selevsek N, Reiter L, Bonner R, Aebersold R (2012) Targeted data extraction of the MS/MS spectra generated by data-independent acquisition: a new concept for consistent and accurate proteome analysis. Mol Cell Proteomics 11: O111 016717.

Grasso CS, Wu YM, Robinson DR, Cao X, Dhanasekaran SM, Khan AP, Quist MJ, Jing X, Lonigro RJ, Brenner JC, Asangani IA, Ateeq B, Chun SY, Siddiqui J, Sam L, Anstett M, Mehra R, Prensner JR, Palanisamy N, Ryslik GA et al. (2012) The mutational landscape of lethal castration-resistant prostate cancer. Nature 487: 239–43.

Guo T, Kouvonen P, Koh CC, Gillet LC, Wolski WE, Rost HL, Rosenberger G, Collins BC, Blum LC, Gillessen S, Joerger M, Jochum W, Aebersold R (2015a) Rapid mass spectrometric conversion of tissue biopsy samples into permanent quantitative digital proteome maps. Nature medicine 21: 407–13.

Guo T, Kouvonen P, Koh CC, Gillet LC, Wolski WE, Rost HL, Rosenberger G, Collins BC, Blum LC, Gillessen S, Joerger M, Jochum W, Aebersold R (2015b) Rapid mass spectrometric conversion of tissue biopsy samples into permanent quantitative digital proteome maps. Nat Med

Haffner MC, Aryee MJ, Toubaji A, Esopi DM, Albadine R, Gurel B, Isaacs WB, Bova GS, Liu W, Xu J, Meeker AK, Netto G, De Marzo AM, Nelson WG, Yegnasubramanian S (2010) Androgen-induced TOP2B-mediated double-strand breaks and prostate cancer gene rearrangements. Nat Genet 42: 668–75.

Haffner MC, Mosbruger T, Esopi DM, Fedor H, Heaphy CM, Walker DA, Adejola N, Gurel M, Hicks J, Meeker AK, Halushka MK, Simons JW, Isaacs WB, De Marzo AM, Nelson WG, Yegnasubramanian S (2013) Tracking the clonal origin of lethal prostate cancer. J Clin Invest 123: 4918–22.

Hayes JH, Barry MJ (2014) Screening for prostate cancer with the prostate-specific antigen test: a review of current evidence. JAMA 311: 1143–9.

Humphrey PA, Moch H, Cubilla AL, Ulbright TM, Reuter VE (2016) The 2016 WHO Classification of Tumours of the Urinary System and Male Genital Organs-Part B: Prostate and Bladder Tumours. Eur Urol 70: 106–119.

Iglesias-Gato D, Wikstrom P, Tyanova S, Lavallee C, Thysell E, Carlsson J, Hagglof C, Cox J, Andren O, Stattin P, Egevad L, Widmark A, Bjartell A, Collins CC, Bergh A, Geiger T, Mann M, Flores-Morales A (2016) The Proteome of Primary Prostate Cancer. Eur Urol 69: 942–52.

Jones S, Chen WD, Parmigiani G, Diehl F, Beerenwinkel N, Antal T, Traulsen A, Nowak MA, Siegel C, Velculescu VE, Kinzler KW, Vogelstein B, Willis J, Markowitz SD (2008) Comparative lesion sequencing provides insights into tumor evolution. Proc Natl Acad Sci U S A 105: 4283–8.

Kamburov A, Stelzl U, Lehrach H, Herwig R (2013) The ConsensusPathDB interaction database: 2013 update. Nucleic Acids Res 41: D793–800

Kristiansen G (2018) Markers of clinical utility in the differential diagnosis and prognosis of prostate cancer. Mod Pathol 31: S143–155

Levine JH, Simonds EF, Bendall SC, Davis KL, Amir el AD, Tadmor MD, Litvin O, Fienberg HG, Jager A, Zunder ER, Finck R, Gedman AL, Radtke I, Downing JR, Pe’er D, Nolan GP (2015) Data-Driven Phenotypic Dissection of AML Reveals Progenitor-like Cells that Correlate with Prognosis. Cell 162: 184–97.

Liu Y, Buil A, Collins BC, Gillet LC, Blum LC, Cheng LY, Vitek O, Mouritsen J, Lachance G, Spector TD, Dermitzakis ET, Aebersold R (2015) Quantitative variability of 342 plasma proteins in a human twin population. Mol Syst Biol 11: 786.

MacLean B, Eng JK, Beavis RC, McIntosh M (2006) General framework for developing and evaluating database scoring algorithms using the TANDEM search engine. Bioinformatics 22: 2830–2.

Magi-Galluzzi C, Xu X, Hlatky L, Hahnfeldt P, Kaplan I, Hsiao P, Chang C, Loda M (1997) Heterogeneity of androgen receptor content in advanced prostate cancer. Mod Pathol 10: 839–45.

Murtaza M, Dawson SJ, Pogrebniak K, Rueda OM, Provenzano E, Grant J, Chin SF, Tsui DW, Marass F, Gale D, Ali HR, Shah P, Contente-Cuomo T, Farahani H, Shumansky K, Kingsbury Z, Humphray S, Bentley D, Shah SP, Wallis M et al. (2015) Multifocal clonal evolution characterized using circulating tumour DNA in a case of metastatic breast cancer. Nat Commun 6: 8760.

Picotti P, Clement-Ziza M, Lam H, Campbell DS, Schmidt A, Deutsch EW, Rost H, Sun Z, Rinner O, Reiter L, Shen Q, Michaelson JJ, Frei A, Alberti S, Kusebauch U, Wollscheid B, Moritz RL, Beyer A, Aebersold R (2013) A complete mass-spectrometric map of the yeast proteome applied to quantitative trait analysis. Nature 494: 266–70.

Plass C, Pfister SM, Lindroth AM, Bogatyrova O, Claus R, Lichter P (2013) Mutations in regulators of the epigenome and their connections to global chromatin patterns in cancer. Nat Rev Genet 14: 765–80.

Powell BS, Lazarev AV, Carlson G, Ivanov AR, Rozak DA (2012) Pressure cycling technology in systems biology. Methods Mol Biol 881: 27–62.

Remmele W, Stegner HE (1987) [Recommendation for uniform definition of an immunoreactive score (IRS) for immunohistochemical estrogen receptor detection (ER-ICA) in breast cancer tissue]. Pathologe 8: 138–40.

Rosenberger G, Koh CC, Guo T, Rost HL, Kouvonen P, Collins BC, Heusel M, Liu Y, Caron E, Vichalkovski A, Faini M, Schubert OT, Faridi P, Ebhardt HA, Matondo M, Lam H, Bader SL, Campbell DS, Deutsch EW, Moritz RL et al. (2014) A repository of assays to quantify 10,000 human proteins by SWATH-MS. Sci Data 1: 140031.

Röst HL, Rosenberger G, Navarro P, Gillet L, Miladinovic SM, Schubert OT, Wolski W, Collins BC, Malmstrom J, Malmstrom L, Aebersold R (2014) OpenSWATH enables automated, targeted analysis of data-independent acquisition MS data. Nat Biotechnol 32: 219–23.

Russnes HG, Navin N, Hicks J, Borresen-Dale AL (2011) Insight into the heterogeneity of breast cancer through next-generation sequencing. J Clin Invest 121: 3810–8.

Schubert OT, Gillet LC, Collins BC, Navarro P, Rosenberger G, Wolski WE, Lam H, Amodei D, Mallick P, MacLean B, Aebersold R (2015) Building high-quality assay libraries for targeted analysis of SWATH MS data. Nat Protoc 10: 426–41.

Shah RB, Bentley J, Jeffery Z, DeMarzo AM (2015) Heterogeneity of PTEN and ERG expression in prostate cancer on core needle biopsies: implications for cancer risk stratification and biomarker sampling. Hum Pathol 46: 698–706.

Sturm M, Bertsch A, Gropl C, Hildebrandt A, Hussong R, Lange E, Pfeifer N, Schulz-Trieglaff O, Zerck A, Reinert K, Kohlbacher O (2008) OpenMS - an open-source software framework for mass spectrometry. BMC Bioinformatics 9: 163.

Tukey JW (1977) Exploratory Data Analysis. Addison-Wesley. ISBN 0-201-07616-0. OCLC 3058187.

Tuxhorn JA, Ayala GE, Smith MJ, Smith VC, Dang TD, Rowley DR (2002) Reactive stroma in human prostate cancer: induction of myofibroblast phenotype and extracellular matrix remodeling. Clin Cancer Res 8: 2912–23.

Uhlen M, Fagerberg L, Hallstrom BM, Lindskog C, Oksvold P, Mardinoglu A, Sivertsson A, Kampf C, Sjostedt E, Asplund A, Olsson I, Edlund K, Lundberg E, Navani S, Szigyarto CA, Odeberg J, Djureinovic D, Takanen JO, Hober S, Alm T et al. (2015) Proteomics. Tissue-based map of the human proteome. Science 347: 1260419.

Umbehr M, Kessler TM, Sulser T, Kristiansen G, Probst N, Steurer J, Bachmann LM (2008) ProCOC: the prostate cancer outcomes cohort study. BMC Urol 8: 9.

Vanhara P, Hampl A, Kozubik A, Soucek K (2012) Growth/differentiation factor-15: prostate cancer suppressor or promoter? Prostate Cancer Prostatic Dis 15: 320–8.

Wettstein MS, Saba K, Umbehr MH, Murtola TJ, Fankhauser CD, Adank JP, Hofmann M, Sulser T, Hermanns T, Moch H, Wild P, Poyet C (2017) Prognostic Role of Preoperative Serum Lipid Levels in Patients Undergoing Radical Prostatectomy for Clinically Localized Prostate Cancer. Prostate 77: 549–556.

Wisniewski JR, Ostasiewicz P, Dus K, Zielinska DF, Gnad F, Mann M (2012) Extensive quantitative remodeling of the proteome between normal colon tissue and adenocarcinoma. Mol Syst Biol 8: 611.

Yachida S, Jones S, Bozic I, Antal T, Leary R, Fu B, Kamiyama M, Hruban RH, Eshleman JR, Nowak MA, Velculescu VE, Kinzler KW, Vogelstein B, Iacobuzio-Donahue CA (2010) Distant metastasis occurs late during the genetic evolution of pancreatic cancer. Nature 467: 1114–7.

Zellweger T, Gunther S, Zlobec I, Savic S, Sauter G, Moch H, Mattarelli G, Eichenberger T, Curschellas E, Rufenacht H, Bachmann A, Gasser TC, Mihatsch MJ, Bubendorf L (2009) Tumour growth fraction measured by immunohistochemical staining of Ki67 is an independent prognostic factor in preoperative prostate biopsies with small-volume or low-grade prostate cancer. Int J Cancer 124: 2116–23.

Zhang B, Wang J, Wang X, Zhu J, Liu Q, Shi Z, Chambers MC, Zimmerman LJ, Shaddox KF, Kim S, Davies SR, Wang S, Wang P, Kinsinger CR, Rivers RC, Rodriguez H, Townsend RR, Ellis MJ, Carr SA, Tabb DL et al. (2014) Proteogenomic characterization of human colon and rectal cancer. Nature 513: 382–7.

Zhang H, Liu T, Zhang Z, Payne SH, Zhang B, McDermott JE, Zhou JY, Petyuk VA, Chen L, Ray D, Sun S, Yang F, Chen L, Wang J, Shah P, Cha SW, Aiyetan P, Woo S, Tian Y, Gritsenko MA et al. (2016) Integrated Proteogenomic Characterization of Human High-Grade Serous Ovarian Cancer. Cell 166: 755–65.

